# Population genetic data of the 21 autosomal STRs included in GlobalFiler^TM^ kit of a population sample from the Kingdom of Bahrain

**DOI:** 10.1101/550400

**Authors:** Noora R. Al-Snan, Safia Messaoudi, Saranya R. Babu, Moiz Bakhiet

**Affiliations:** Department of Molecular Medicine, College of Medical and Medicine Sciences, Arabian Gulf University, Kingdom of Bahrain; Forensic Science Laboratory, Directorate of Forensic Science, General Directorate of Criminal Investigation and Forensic Science, Ministry of Interior, Kingdom of Bahrain; Forensic Biology Department, College of Forensic Sciences, Naif Arab University for Security Sciences, Riyadh, Saudi Arabia

**Keywords:** Bahraini population, GlobalFiler(^TM^) PCR kit, Forensic parameters, Population genetics

## Abstract

**Introduction:** Bahrain’s population consists mainly of Arabs, Baharna and Persians leading Bahrain to become ethnically diverse. The exploration of the ethnic origin and genetic structure within the Bahraini population is fundamental mainly in the field of population genetics and forensic science.

**Aim:** The purpose of the study was to investigate and conduct genetic studies in the population of Bahrain to assist in the interpretation of DNA-based forensic evidence and in the construction of appropriate databases.

**Materials and Methods:** 24 short-tandem repeats in the GlobalFiler™ PCR Amplification kit including 21 autosomal STR loci and three gender determination loci were amplified to characterize different genetic and forensic population parameters in a cohort of 543 Bahraini unrelated healthy men. Samples were collected during the year 2017.

**Results:** The genotyping of the 21 autosomal STRs showed that most loci were in Hardy-Weinberg Equilibrium (HWE) except for three markers; D3S1358, D19S433 and D5S818 which showed deviation from HWE. We also found out no significant deviations from LD between pairwise STR loci in Bahraini population except when plotting for D3S1358-CSF1PO, CSF1PO-SE33, D19S433-D12S391, FGA-D2S1338, FGA-SE33, FGA-D7S820 and D7S820-SE33. The SE33 locus was the most polymorphic for the studied population and THO1 locus was the less polymorphic. The Allele 8 in TPOX scored the highest allele frequency of 0.496. The SE33 locus showed the highest power of discrimination (PD) in Bahraini population, whereas TPOX showed the lowest PD value. The 21 autosomal STRs showed a value of combined match probability (CMP) equal to 4.5633^E-27^, and a combined power of discrimination (CPD) of 99.99999999%. Off-ladders and tri-allelic variants were observed in various samples at D12S391, SE33 and D22S1045 loci.

**Conclusion:** Our study indicated that the twenty-one autosomal STRs are highly polymorphic in the Bahraini population and can be used as a powerful tool in forensics and population genetic analyses including paternity testing and familial DNA searching.

**Author Summary:** Kingdom of Bahrain is a country of 33 islands located in the Arabian Peninsula. The location of Bahrain had affected the diversity of its population, which is mainly divided into four main ethnic groups: Arabs, Baharna and Persians. Genetic studies on Bahraini population are very limited and little has been done to characterize population structure within Kingdom of Bahrain. Here, we used 21 autosomal STRs included in the GlobalFiler™ Amplification Kit to amplify DNA from 543 non-related males from Bahraini population. We conducted statistical analysis using two main different software such as STRAF and GenAlEx. Different forensic and population parameters were obtained to characterize Bahraini population. Some of the significant results obtained were the following: most of the loci were in Hardy-Weinberg Equilibrium, the most polymorphic and informative marker was SE33. Allele 8 in TPOX presented the highest allele frequency for the studied population. We also found out some of the rare variants which were recorded in STRbase website. Bahraini population was correspondingly compared to the genetic structure of the region. Our study indicated the usefulness of the 21 autosomal STRs in the GlobalFiler ™ in establishing databases, analyzing paternity and reviewing DNA-based evidences.

## 1. Introduction

Kingdom of Bahrain is a small archipelago consisting of 33 islands, only the five largest are inhabited. These islands are Bahrain, Muharraq, Umm and Nasan and Sitra. Bahrain is positioned in the Arabian Gulf. To the southeast of Bahrain is the State of Qatar, and to its west lies the Kingdom of Saudi Arabia, with which it is connected by a 25-kilometer causeway. To the north and east of Bahrain lies the Islamic Republic of Iran **(1).**

Bahrain is one of the most densely populated countries in the world, with a total landmass of 760 square kilometers. Mid-2014, estimates of Bahrain’s population stood at 1,314,562 persons. Of these, 568,399 are Bahraini citizens (46%) and 666,172 are expatriates (54%) **(2).**

Standing between the most substantial focal points of the ancient world – the Far East, the Indus Valley, Fertile Crescent, the Red Sea and the Coast of East Africa **(3)**, trade goods from the Persian Gulf made its way into Europe through Antioch **(4).** This made Bahrain an important port city, a metropolitan hub where different cultures met **(5).**

Because of the geographic location of Bahrain, the diversity of the population had been affected. This could be explained by the migration flows from several areas regionally, and eventually internationally **(6).** Iranians and migrants of Iranian heritage constituted the largest groups of migrants who were Muslim and ethnically not Arab **(7)**. Indian and Iranian migration boomed in the early and mid-20th century, as the Bahrain Petroleum Company sought a workforce for the oil that was discovered in the island **(8).**

Population is mainly divided into four main ethnic groups: Arabs, Baharna and Persians (Huwala and Ajam) **(4, 9, 10).** This geographical and social organization might be expected to have an effect on patterns of a genetic diversity **(11).**

Genetic studies on Bahrain to date are very limited and knowledge of any such structure is important in the interpretation of the significance of DNA-based forensic evidence and in the construction of appropriate databases. This present study is the first to characterize genetically the Bahraini population, using Globalfiler™ amplification kit. Twenty-four autosomal short-tandem repeats (STRs) in GlobalFiler™ PCR Amplification kit (Thermo Fisher Scientific, Inc., Waltham, MA, USA) were studied to characterize different forensic and genetic population parameters in 545 Bahraini males. The 6-dye GlobalFiler™ PCR Amplification kit (Thermo Fisher Scientific, Inc., Waltham, MA, USA) was designed to incorporate 21 commonly used autosomal STR loci (D8S1179, D21S11, D7S820, CSF1PO, D3S1358, TH01, D13S317, D16S539, D2S1338, D19S433, VWA, TPOX, D18S51, D5S818, FGA, D12S391, D1S1656, D2S441, D10S1248, D22S1045 and SE33) and three gender determination loci (Amelogenin, Yindel and DYS391) which have been proven to provide reliable DNA typing results and enhance the power of discrimination (PD).

## 2. Results and discussions

### 2.1. Hardy Weinberg Equation (HWE) and Linkage Disequilibrium (LD)

In the present study no significant deviation from HWE was observed (p> 0.05) except for three markers; D3S1358, D19S433 and D5S818 **(Table 1).** This observed deviation could be a result of the diversity of the Bahraini population or caused by high polymorphism at the same loci investigated loci. This observation are likely to reflect the high level of inbreeding with consanguinity rates in Bahrain, with intra-familial unions accounting for 20–50% of all marriages compared to other Arab countries **(12)**. D22S1015, SE33 and D21S11 loci also revealed evidence of a rare variant and off-ladders **(Figure 1).**

**Figure 1:**
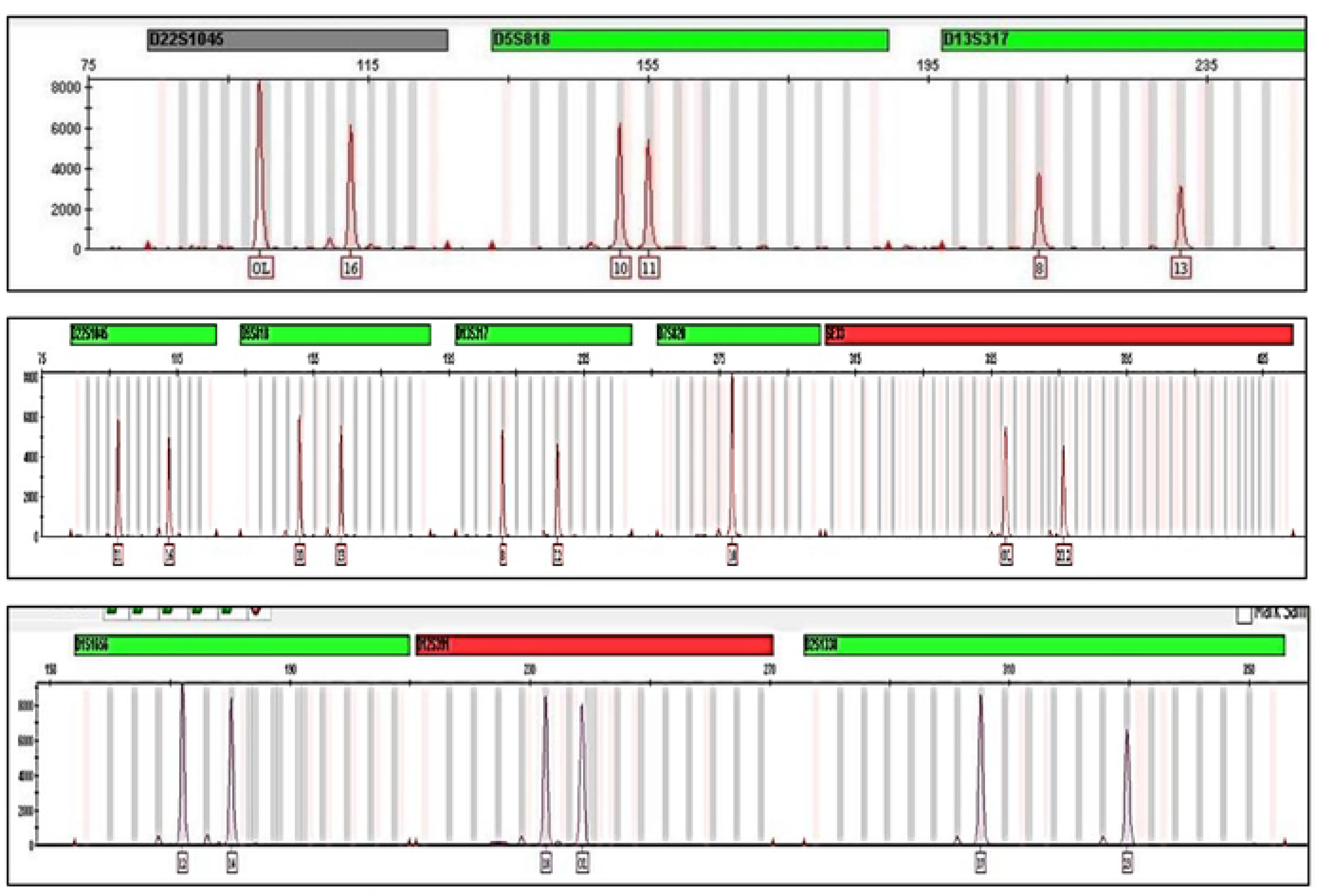
Off-ladder cases observed in D22S0145, SE33 and D12S391.

The study also showed no significant deviation from LD between pairwise STR loci after Bonferroni’s correction (p>0.000092) in Bahraini population except for the following loci; D3S1358-CSF1PO, CSF1PO-SE33, D19S433-D12S391, FGA-D2S1338, FGA-SE33, FGA-D7S820 and D7S820-SE33 when plotted. The highest pairwise LD was 1.00 when plotting CSF1PO-D19S433, D21S11-FGA and FGA-D1S1656. The marker D22S1045 did not show any probability. This lack of probability correlated with the off-ladder cases observed in D22S1045 and which may be the reason for the null probability value.

### 2.2. Allele frequencies and Forensic parameters

In the studied population, the number of allele (Na) per locus was ranged from 7 for markers D16S539, TPOX and THO1 to 48 for SE33, the mean number of alleles per locus was 14, and a total number of alleles observed was 288. The most polymorphic locus was SE33 **(Table 1).**

The probability that two randomly chosen person have the same unspecified genotype at a locus is the sum squares of the frequencies of all genotypes at that locus. Some alleles show very high frequencies in the Bahraini population; allele 8 in locus TPOX scored the highest frequencies of 0.496 followed by allele 15 in D22S1045 with frequency of 0.417 and the lowest allele frequency was 0.00092 for 35 different alleles. **(Table 1).**

Generally, the polymorphism degree of a specific locus can be measured by two distinct parameters – the heterozygosity and the Polymorphism Information Content (PIC) of this locus. We have found out that the observed heterozygosity (Ho) was ranged from 67% for locus TPOX to 92% for locus SE33. **(Table 1).**

The Polymorphism Information Content (PIC) values for all STR loci were highly informative (PIC≥0.6) with an average of 78.3%.

The means for (Na) and (He) designate the high levels of genetic diversity in the population studied. These high informative values support the heterozygosity values indicating the high degree of genetic polymorphism.

The random matching probability (PM) was ranged from 0.006 for SE33 to 0.156 for TPOX. The Power of exclusion (PE) was ranged from 0.384 for locus TPOX to 0.838 for locus SE33. The SE33 locus showed the greatest (PD) in Bahraini population, whereas TPOX showed the lowest. The higher the discrimination power of a locus, the more efficient it is in discriminating between members of the population **(Table 1).**

The PD values for most of the tested loci was above 0.9; the highest observed at SE33 with 0.994 and the least at TPOX with 0.844. The combined power of discrimination (CPD) and combined matching probability (CMP) for all the 21 STR loci were 99.999999% and 4.5633^E-27^ respectively. The PD in correlation with PM supports the high degree of polymorphism between Bahraini individuals.

### 2.3. Genetic structure in the region

We have compared Bahraini population data to the nearest available populations using the accessible loci. It is shown that the Bahraini population shares similar results with the study conducted of Saudi Arabia and UAE populations using the GlobalFiler™ STR loci **(13, 14)**. As the above-mentioned populations share the most informative and polymorphic locus is SE33 and the least informative locus is TPOX. The least polymorphic was locus D16S539 for UAE population **(14)** whereas THO1 for both Bahraini and Saudi Arabian populations **(13).** Allele 8 in locus TPOX scored the highest frequency for Bahraini, Kuwaiti, Saudi Arabian, Iraqi, Egyptian and Iranian populations **(13, 15)** whereas the highest frequency for Indian and Bangladeshi populations is allele 12 in CSF1PO **(15).** As expected, the diversity between the data obtained in this study compared to the neighboring data populations varies, as the populations become more geographically separated.

Once more studies of Arab populations in the region become accessible, it may be more probable to develop a greater understanding of the genetic associations between the different populations for the Arabian Peninsula.

### 2.4. Rare variants, off-ladder and null alleles

Different samples showed off ladder (OL) in 10 various cases; two allelic ladder variants were detected at the D12S391, Sample#5 indicated OL (18,OL) in 238.69 bp and sample#511 showed OL (OL,21) in 238.64 bp. Two allelic ladder variants were detected at the SE33, Sample#288 indicated OL in (OL,23.2) in 320.61 bp and Sample#538 with OL (OL,21.2) in 359.20 bp. Six allelic ladder variants were detected in D22S1045, sample#309 showed OL (OL,17) in 99.44 bp, sample#331 showed OL (OL,15) in 99.51 bp, sample#487 showed another OL (OL,14) in 99.45 bp, sample#516 indicated an OL (OL,16) in 99.44 bp, sample#524 showed OL (OL,14) in 99.45 bp and sample #549 showed OL (OL,16) in 99.44 bp **(Figure 1).** As for the tri-allelic patterns, sample#180 showed three variants in D21S11 (30,31.2,32.2) with sizes of 207.67 bp, 213.64 bp and 217.78 bp **(Figure 2A)** respectively and it was not observed and reported in STRbase (http://strbase.nist.gov/index.htm) **(16).** Sample# 520 showed 3 variants in D2S441 (10,11,12) with sizes of 85.03 bp, 89.15 bp and 93.30 bp respectively (**Figure 2B)**. Whereas the adjacent locus D19S433 was of homozygous allele (13,13) and it was observed and previously reported in STRbase (http://strbase.nist.gov/index.htm) **(16).**

**Figure 2:**
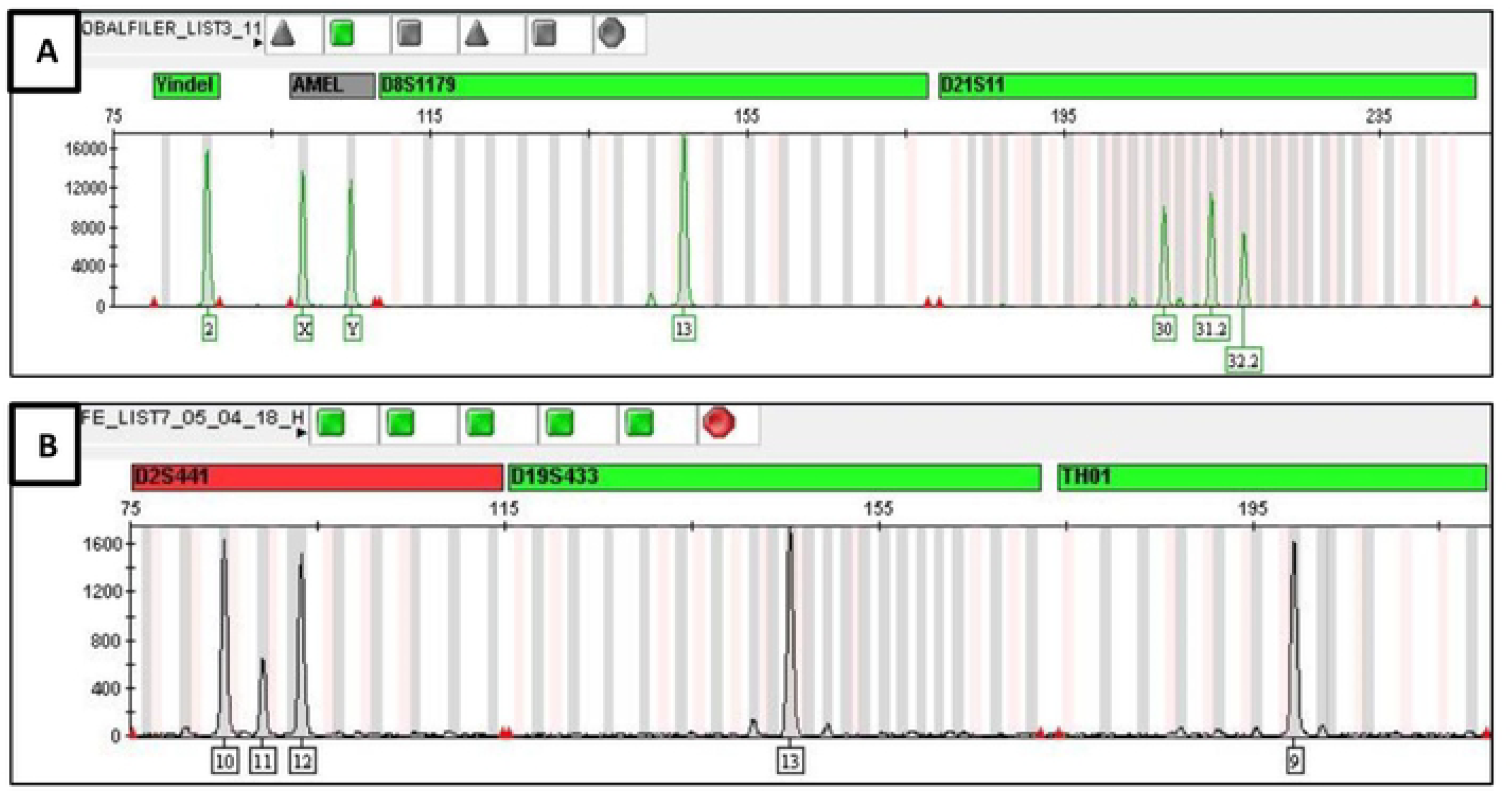
Two electropherograms (A & B) indicating the tri allelic patterns.

## 3. Materials and methods

### 3.1. Sample Collection

Five hundred and forty-three (543) blood samples were collected on Nucleic-Cards™ (Copan, Italy) from non-relatives’ Bahraini males. In each case, males with ancestry (to the level of paternal grandfather) from four different geographical subdivisions of the country (Capital Governorate, Muharraq Governorate, Northern Governorate and Southern Governorate) were sampled. Ethical review for analysis was provided by the Research and Research Ethics Committee (RREC) (E007-PI-10/17) in the Arabian Gulf University. All participants provided informed consent.

### 3.2. DNA amplification and fragment detection

DNAs were punched and amplified from Nucleic-Cards™ (Copan, Italy) blood-spot samples using a fully automated workstation, starting from 1.2-mm diameter punches produced using the easyPunch STARlet system (Hamilton, Switzerland).

The extracted DNA samples were directly amplified using GlobalFiler™ (Thermo Fisher Scientific, Inc., Waltham, MA, USA) according to manufacturer’s recommendation. A total of 24 loci were amplified, including 21 autosomal STR loci and three gender determination loci.

The PCR products were separated by capillary electrophoresis in an ABI 3500xl Genetic Analyzer (Thermo Fisher Scientific Company, Carlsbad, USA) with reference to the LIZ600 size standard v2 (Thermo Fisher Scientific, Inc., Waltham, MA, USA) and the GeneMapper® ID-X Software v1.4 (Thermo Fisher Scientific, Inc., Waltham, MA, USA) was used for genotype assignment. DNA typing and assignment of nomenclature were based on the ISFG recommendations

### 3.3. Statistical analysis

Allele frequencies, Minor allele frequencies (MAF) and different parameters of forensic efficiency—such as power of discrimination (PD), random matching probability (PM), power of exclusion (PE), polymorphism information content (PIC), typical paternity index (TPI), and heterozygosity (He)—were estimated for each locus using GenAlEx software V.6.503 **(17).** Fisher’s exact tests to evaluate the Hardy–Weinberg equilibrium (HWE) by locus and linkage disequilibrium (LD) between pair of loci were estimated with STRAF - A convenient online tool for STR data evaluation in forensic genetics **(18).**

## 4. Conclusions

In conclusion, we have reported the allele frequencies and forensic statistical parameters of the GlobalFiler™ STR loci in Bahraini population to be indicated in literature for the first time. The polymorphism of the 21 autosomal markers observed in this study such as SE33 marker indicates its usefulness for paternity testing, forensics and familial DNA searching in the population of Bahrain.

Overall, these parameters indicated the general utility of this STR loci panel for forensic personal identification and paternity testing in the Bahraini population, thereby further confirming of its efficacy for forensic practice also in Bahraini sub-populations and other populations’ genetics and diversity studies.

## Conflict of interest

The authors declare that they have no conflict of interest

## Acknowledgments

We would like to thank the authorities in General Directorate of Criminal Investigation and forensic Science in Bahrain, namely Mr. Abdulaziz Mayoof Al-Rumaihi and Mr. Mohammed Abdulla Ghayyath for allowing us to utilize the Bahrain forensic Science Laboratory. Also, many thanks to Latifa Ahmed and Sabah Nazir for their technical support. This research did not receive any specific grant from funding agencies in the public, commercial, or not-for-profit sectors.

